# NIAGADS Alzheimer’s GenomicsDB: A resource for exploring Alzheimer’s Disease genetic and genomic knowledge

**DOI:** 10.1101/2020.09.23.310276

**Authors:** Emily Greenfest-Allen, Otto Valladares, Pavel P. Kuksa, Prabhakaran Gangadharan, Wan-Ping Lee, Jeffrey Cifello, Zivadin Katanic, Amanda B. Kuzma, Nicholas Wheeler, William S. Bush, Yuk Yee Leung, Gerard Schellenberg, Christian J. Stoeckert, Li-San Wang

**Affiliations:** Penn Neurodegeneration Genomics Center, Perelman School of Medicine, University of Pennsylvania, Philadelphia, PA 19104, USA; Institute for Biomedical Informatics, Perelman School of Medicine, University of Pennsylvania, Philadelphia, PA 19104, USA; Department of Pathology and Laboratory Medicine, Perelman School of Medicine, University of Pennsylvania, Philadelphia, PA 19104, USA; Department of Genetics, Perelman School of Medicine, University of Pennsylvania, Philadelphia, PA 19104, USA; Cleveland Institute for Computational Biology, Department of Population and Quantitative Health Sciences, Case Western Reserve University, Cleveland, OH, 44106, USA

## Abstract

**INTRODUCTION:** The NIAGADS Alzheimer’s Genomics Database (GenomicsDB) is a public knowledgebase of Alzheimer’s disease (AD) genetic datasets and genomic annotations.

**METHODS:** It uses a custom systems architecture to adopt and enforce rigorous standards that facilitate harmonization of AD-relevant GWAS summary statistics datasets with functional annotations, including a database of >230 million annotated variants from the AD Sequencing Project’s joint-calling efforts.

**RESULTS:** The knowledgebase generates genome browser tracks and interactive compiled from harmonized datasets and annotations in the underlying database. These facilitate data sharing and discovery, by contextualizing AD-risk associations in a broader functional genomic context or summarizing them in the context of functionally annotated genes and variants.

**DISCUSSION:** Created to make AD-genetics knowledge more accessible to AD-researchers, the GenomicsDB shares annotated AD-relevant summary statistics datasets via a web interface designed to guide users unfamiliar with genetic data in not only exploring, but also interpreting this ever-growing volume of data.

## 1 Introduction

Alzheimer’s disease (AD) is a progressive neurodegenerative disorder that affects approximately 6.7 million Americans aged 65 or older as of 2023 [1], is effectively untreatable, and invariably progresses to complete incapacitation and death 10 or more years after onset. Our current understanding of genetic risk for AD has resulted mainly from ongoing massive genotyping and sequencing efforts carried out by groups such as the Alzheimer’s Disease Genetics Consortium (ADGC), the International Genomics of Alzheimer’s Project (IGAP), and the Alzheimer’s Disease Sequencing Project (ADSP). Large-scale genome wide association studies (GWAS) and GWAS-derived meta-analyses have been performed by each of these groups [2–5]. The results of these and other recent AD-GWAS analyses have greatly expanded the known list of AD-risk associated variants, identifying as many as 75 AD risk-associated loci to date [6]. Most of these loci lie within non-coding regions of the genome, necessitating downstream analyses that annotate variants by known pathogenicity and weigh association strength against proximal genomic features to elucidate causal variants, impacted regulatory elements, and potential therapeutic targets for mitigating AD/ADRD pathologies. Such analyses typically integrate GWAS summary statistics (containing at a minimum *p*-values reflecting association strength) with a variety of genomic annotations, ranging from genomic features (e.g., Ensembl genes and transcripts [7]) to regulatory elements and epigenomic or transcriptomic annotations in “disease-relevant” tissues or cell populations (e.g., brain, cerebrospinal fluid, microglia and other neuronal cells) selectively pulled from large-scale projects such as the Encyclopedia of DNA Elements (ENCODE) [8,9], the Functional Annotation of the Mouse/Mammalian Genome (FANTOM5) enhancer atlas [10], and the Genotype-Tissue Expression Portal (GTEx) [11]. Consequently, characterizing the potential regulatory impact or identifying target “effector” genes of a GWAS locus requires substantial bioinformatics effort as data from disparate resources need to be standardized to facilitate harmonization and enable accurate (and efficient) mapping of genomic annotations to the GWAS results, often on genome-wide scales.

Results from ADGC, ADSP and IGAP GWAS and other AD-relevant GWAS studies, including their summary statistics, are deposited at the National Institute of Aging (NIA) Genetics of Alzheimer’s Disease Data Storage Site (NIAGADS). NIAGADS is a NIA-designated essential national infrastructure, providing a one-stop access portal for Alzheimer’s disease ‘omics datasets [12]. Investigators can gain access to these datasets through a qualified data access request (DAR) process. NIAGADS also makes *p*-values (see **section 2.2.6** for details) from >100 GWAS summary statistics and meta-analysis datasets (e.g., single variant and rare variant aggregation tests of association) available for public download (no DAR required). Most were generated for the GRCh37/hg19 genome build, but newer GRCh38/hg38 datasets are now available at NIAGADS, with more incoming.

The NIAGADS collection of GWAS summary statistics datasets is unique, in that it provides a disease-centric resource for AD researchers. However, AD-relevant datasets are also available from large-scale manually curated disease-agnostic collections such as the MRC-IEU Open GWAS Project (OpenGWAS) [13] and the NHGRI-EBI GWAS Catalog [14]. OpenGWAS currently includes 12 GRCh37/hg19 aligned AD-GWAS summary statistics datasets, a quarter of which are derived from ADGC or IGAP-studies deposited at NIAGADS. The GWAS Catalog has a larger offering, listing 15 publications associated with the term “Alzheimer’s disease” that have full sets of AD-related summary statistics that can be downloaded by the public (n = 38 datasets in total, some of which overlap with OpenGWAS and NIAGADS offerings on both genome builds).

Despite the general availability of these GWAS summary statistics datasets, there are three sizeable challenges that preclude the AD-research community from taking full advantage of this large and ever-increasing volume of data. These are 1) the systematic integration and normalization of the summary datasets themselves, which can vary in aspects of the study design (e.g., number and selection criteria for samples based on available phenotypes, statistical methods and adjustments) and in format and contents of the files (e.g., labelling of risk-associated variants, which statistics are included); 2) the computational overhead and bioinformatics expertise involved in the downstream analyses that are necessary to place GWAS loci in a broader genomic context and elucidate causal variants and their potential molecular or functional impact; and 3) the ability to disseminate the contextualized association data in a standardized, organized, easily accessed, and searchable format to so that it can be a resource for all AD-researchers (i.e., molecular biologists and clinicians, as well as bioinformaticians).

Here, we describe the NIAGADS Alzheimer’s Genomics Database (GenomicsDB), which was developed in collaboration with the ADGC and ADSP to help overcome these hurdles and make the summary statistics datasets deposited at NIAGADS more accessible to the AD-research community. The GenomicsDB is a user-friendly web-knowledgebase that disseminates annotated AD-relevant GWAS summary statistics datasets, facilitates real-time data mining, and provides summaries of association results in the context of functionally annotated genes and variants.

## 2 Methods

### 2.1 Database infrastructure

An overview of the NIAGADS GenomicsDB systems architecture is provided in **Figure 1**. The GenomicsDB is powered by a PostgreSQL relational database system that has been optimized for parallel big data querying, allowing for efficient real-time data mining. Data are organized using the modular Genomics Unified Schema version 4 (GUS4), designed for scalable integration and dissemination of large-scale ‘omics datasets. Loading of all data is managed by the GUS4 application layer (https://github.com/VEuPathDB/GusAppFramework), which ensures the data integrity and accuracy of data integration. All data preprocessing scripts and NIAGADS customizations to the GUS4 schema and application framework made to harmonize, store, and optimize retrieval of genomic features, annotations, and summary statistics datasets are available on GitHub (https://github.com/NIAGADS/GenomicsDBData/).

**Figure 1.**
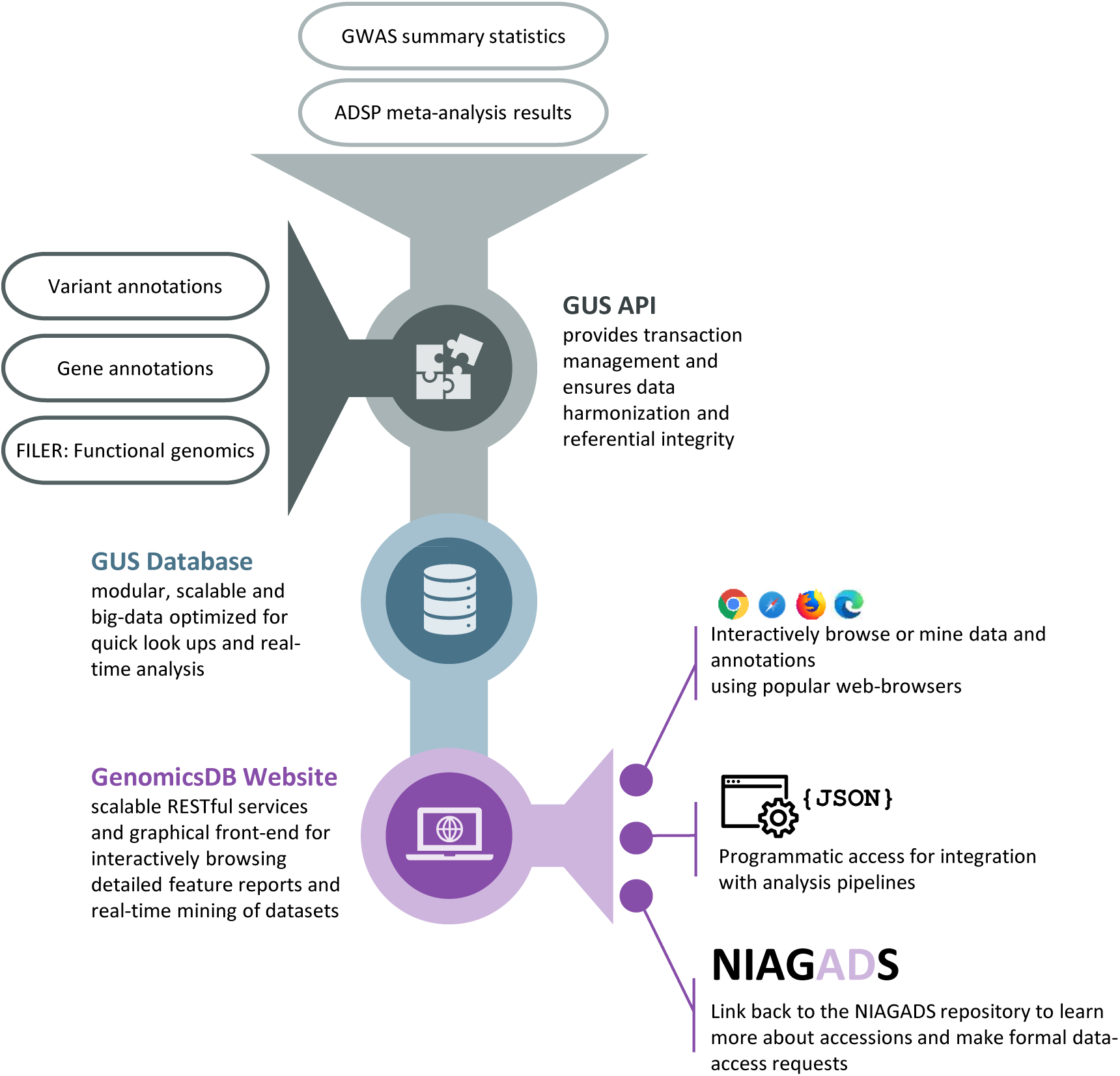
Overview of the GenomicsDB systems architecture. Dataset harmonization and mapping to functional genomics tracks and variant or gene annotations is managed by the GUS application layer. Data are stored in a big data optimized PostgreSQL relational database and organized using the GUS schema (see **section 2.1**), which was designed for scalable integration of large-scale ‘omics datasets. The web development kit used to generate the GenomicsDB website provides RESTful services to efficiently query data from the database that are tied to a graphical front end for interactive data exploration. The same services can also be used to programmatically query the database, allowing integration with data analysis pipelines. The GenomicsDB website supplies quick links back to the NIAGADS repository to facilitate formal data-access requests.

### 2.2 Data harmonization and annotation

In the GenomicsDB, data harmonization depends on two main elements. First variant and gene features are assigned unique identifiers following established conventions that will persist across future versions of the knowledgebase (see **sections 2.2.4** and **2.2.5**, respectively, for details). Adopting standardized feature identifiers is essential for automatic discovery that facilitates large-scale data harmonization. Doing so allows NIAGADS to identify and link equivalent features across disparate annotation resources and in turn also lets third-party resources easily generate permalinks back to GenomicsDB records using templated URLs. Second, indexing genomic locations permits fast retrieval of proximal, overlapping, and co-located elements.

#### 2.2.1 Indexing of genomic features and annotations

All genomic features (incl. genes, gene sub features, and variants) and annotations in the GenomicsDB are sorted and indexed on genomic location using a bin indexing system modeled after that used by the UCSC Genome Browser [15,16]. Each chromosome is split into a set of evenly spaced, nested bins of increasingly finer resolution. During load, features are assigned to the smallest enclosing bin, which are represented in the database as *ltree* objects, a PostgreSQL data type that stores hierarchical data in string dot-path notation to reduce storage overhead, but leverages tree traversal algorithms for fast data querying and retrieval [17,18]. GenomicsDB queries for feature or annotation overlaps with any region of interest assign the queried region to the minimally enclosing bin and then use simple comparison operators to find all annotations within the assigned bin or any parent or nested child bins that overlap the query region.

#### 2.2.2 ADSP variant annotations

As part of their sequencing effort, the ADSP developed an annotation pipeline that builds on Ensembl’s VEP software [19] to efficiently annotate variants and rank potential variant consequences according to the severity of the predicted effect (such as codon changes, loss of function, and potential deleteriousness) [20,21]. Elements of the ADSP annotation pipeline have been integrated with data preprocessing scripts for the GenomicsDB, allowing non-ADSP variants in the knowledgebase to be annotated according to ADSP conventions. ADSP annotations applied to GenomicsDB variants include the ranked variant consequences, allele frequencies, and impacted genes, transcripts, and regulatory elements. Super population allele frequency data are pulled from 1000 Genomes [22] (GRCh37/hg19: phase 3 version 1 [11 May 2011]; GRCh38/hg38: 1000 Genomes 30x [23]), ExAC (GRCh37/hg19 only) [24], the Genome Aggregation Database (gnomAD, https://gnomad.broadinstitute.org/) [25], and the ALFA project (GRCh38/hg38 only) [26]. Allele frequencies from NIAGADS GWAS analyses, ADSP variant calling, and ADSP meta-analyses are restricted and only accessible via formal data access requests to NIAGADS. The pipeline also extracts Combined Annotation Dependent Depletion (CADD) scores [27,28] from the CADD v1.6 release for both genome builds, which quantify and rank potential variant deleteriousness for single nucleotide polymorphisms (SNPs) and short-indels.

#### 2.2.3 Linkage disequilibrium

Linkage-disequilibrium (LD) structure around annotated variants for 1000 Genomes super populations (GRCh37/hg19: phase 3 version 1 [11 May 2011]; GRCh38/hg38: 1000 Genomes 30x [23]) were estimated using PLINK v1.90b2i 64-bit [29]. Only LD-scores meeting a correlation threshold of r^2^ ≥ 0.2 are stored in the database. Available in the GRCh38/hg38 GenomicsDB are LD-estimates for the European (non-Hispanic white) subset of the ADSP R3 17k whole genome sequencing (WGS) samples. These were calculated using emeraLD [30]. Updated LD estimates for ADSP populations will be made available along with future releases of ADSP variants called from WGS. Comparable ADSP WGS results are not available for GRCh37/hg19; no LD for ADSP populations is available for that genome build.

#### 2.2.4 Variant records

SNPs and short-indels are uniquely identified in the GenomicsDB by chromosomal coordinates and allelic variant (chr:position:ref_allele:alt_allele). This allows accurate mapping of risk-association statistics to ADSP variants (see below) and to external variant records in annotation resources such as dbSNP (https://www.ncbi.nlm.nih.gov/snp/) [31], gnomAD, and GTEx (e.g., expression quantitative trait loci, or eQTLs).

All standardized and annotated variants are stored in the PostgreSQL database, creating an in-house reference set against which incoming variants from GWAS summary statistics datasets or third-party annotations can be compared. Approximately 1 billion variants from dbSNP have been annotated using the ADSP annotation pipeline (see **section 2.2.2**) and provide the foundation for this reference set. Identifiers for variants called and passing quality control checks (QC) from ADSP sequencing efforts (ADSP variants) are then standardized and mapped against the dbSNP reference. Found records passing ADSP QC are flagged and updated to include the QC status for the ADSP release; unmatched records are run through the integrated ADSP annotation pipeline and added to the reference variant set. For GRCh37/hg19, the ADSP reference variants are SNPs and short-indels identified during the ADSP Discovery Phase WGS and whole exome sequencing (WES) efforts [20]. For GRCh38/hg38, ADSP reference variants are from a quality checked joint-genotype calling of 16,906 whole-genomes (R3 17k WGS; NIAGADS ACCN: NG00067.v5). Reference variants will be updated or added as additional ADSP join-genotype calls are released to the public.

Risk-associated variants from GWAS summary statistics datasets and trait-associations pulled from the NHGRI-EBI GWAS Catalog are likewise mapped against the set of reference variants. New variant records are annotated and added to the reference when a “novel” (unmatched) variant is found, and a flag is added to the record if the variant has genome-wide significance for AD or an AD-related trait in the association analysis (*p* ≤ 5e^-8^; adjusted to account for false positives due to testing associations of millions of variants simultaneously). The code for standardizing, annotating, and indexing variants comprising this Annotated Reference Variant Database (AnnotatedVDB) is available on GitHub (https://github.com/NIAGADS/AnnotatedVDB).

#### 2.2.5 Gene records

Gene and transcript models are currently pulled from the GENCODE Release 19 (GRCh37/hg19) and GENCODE Release 36 (GRCh38/hg38) reference gene annotation [32]; gene annotations are updated as needed to keep them current. GenomicsDB gene records are uniquely identified by their Ensembl identifiers, as are any sub features (e.g., transcripts, exons) and protein products. Alternative gene identifiers (incl. NCBI Gene IDs, UniProtKB IDs, UCSC Gene IDs) are mapped to Ensembl IDs via the UniProtKB (https://www.uniprot.org/) [33] ID mapping file for human genes; additional standard gene nomenclature is imported from the HUGO Gene Nomenclature Committee at the European Bioinformatics Institute (HGNC) [34]. This includes both official gene symbols and names, as well as homologs in model organisms and identifiers in clinical knowledgebases such as the Online Mendelian Inheritance in Man (OMIM) database (https://omim.org/) [35,36].

These mappings are used to link GenomicsDB records to external resources, as well as harmonize third-party gene annotations not typically mapped to Ensembl IDs, such as pathway and Gene Ontology (GO) associations. Gene membership in molecular and metabolic pathways is obtained from the Kyoto Encyclopedia of Genes and Genomes (KEGG) (https://www.genome.jp/kegg/) [37] and Reactome (https://reactome.org/) [38]. Annotations of the functions of genes and gene products are taken from packaged releases of the Gene Ontology (http://geneontology.org) and GO-gene associations [39] and are updated yearly.

#### 2.2.6 GWAS summary statistics dataset records

GWAS summary statistics datasets deposited at NIAGADS are added to the GenomicsDB as they become publicly available via publication or permission of the submitting researchers. These include studies that focus specifically on AD, as well as those on AD biomarkers and related neuropathologies. An up-to-date listing of the available summary statistics datasets is provided by the GenomicsDB Dataset Browser (https://www.niagads.org/genomics/app/record/dataset/accessions; see also **Fig. 3**).

Prior to loading in the database, metadata is compiled for each dataset to capture provenance, phenotypes (e.g., disease state, neuropathology, population, APOE genotype, etc.), and relevant elements of the study design (e.g., family or case control study, sample size, statistical covariates). Phenotypes are standardized using controlled vocabularies pulled from OBO Foundry ontologies [40,41] to facilitate searches for related datasets and harmonization of trait associations with curated catalogs of GWAS association results (e.g., NHGRI-EBI GWAS Catalog). During the data loading process, variant representations are standardized, indexed, and annotated as described in previous sections. This enables fast lookups and simplifies harmonization with third-party annotations.

To ensure the privacy of personal health information, the NIAGADS GenomicsDB website only makes *p*-values from the summary statistics available for browsing (on dataset, gene, and variant reports and as genome browser tracks) and analysis. Access to the full summary statistics is restricted as under some conditions (e.g., large GWAS sample sizes or family-based studies), some values (e.g., genome-wide allele frequencies) may be personally identifiable [42,43]. The full summary statistics and corresponding GWAS or sequencing results can be obtained via formal data-access requests made to NIAGADS. All datasets are properly credited to the submitting researchers or sequencing project.

The GRCh38/hg38 version of the GenomicsDB provides summary statistics that have been lifted over from GRCh37/hg19. If the original dataset identifies variants only by dbSNP refSNP IDs, GRCh38/hg38 coordinates are determined by mapping the refSNP identifier against the GenomicsDB variant reference set (described in **section 2.2.4**), with checks made for deprecated or synonymized rsIDs. In all other cases a two-stage lift over process is applied. A bed file is generated containing all variant features and their positional information, which is then run through the UCSC Genome Browser liftOver script [15,44]. Any unmapped or uncertain mappings are then submitted to the NCBI coordinate remapping service (Remap; https://www.ncbi.nlm.nih.gov/genome/tools/remap). Any features left unmapped are dropped, as well as long-indels, which are not lifted over from GRCh37/hg19 to GRCh38/hg38 as there is lower confidence that the full sequence is conserved or that the allele sequences will still be valid.

### 2.3 Website design and organization

The NIAGADS GenomicsDB is powered by an open-source database system and web-development kit (WDK; https://github.com/VEuPathDB/WDK) developed and successfully deployed by the Eukaryotic Pathogen, Vector and Host Informatics (VEuPathDB) Bioinformatics Resource Center [45]. The VEuPathDB WDK provides a query engine that ties the database system to the website via an easily extensible XML data model. The data model is used to automatically generate and organize searches, search results, and reports, with concepts and data organized by topics from the EMBRACE Data And Methods (EDAM) ontology, which defines a comprehensive set of concepts that are prevalent within bioinformatics [46]. This facilitates updates of third-party data and rapid integration of new datasets as they become publicly available.

The WDK also provides a framework for lightweight Java/Jersey representational state transfer (REST) services for data querying. The framework allows search results and reports to be returned in multiple file formats (e.g., delimited-text, XML, and JSON) in addition to browsable, interactive web pages. The user interface is powered by a React/JS framework that queries the REST services and dynamically updates website content and displays depending on user choices. All code used to generate the website, JavaScript genomics visualizations. are available on GitHub (https://github.com/NIAGADS/GenomicsDBWebsite).

A combination of in-house JavaScript genomics visualizations and third-party visualization toolkits are used to provide graphical interfaces for browsing, summarizing, and mining datasets and annotations in gene and variant reports. This includes interactive *LocusZoom.js* [47,48] plots for viewing risk-associated loci in the context of the local LD-structure. Custom data adapters have been written for *LocusZoom.js* that leverage the GenomicsDB REST services to pull summary statistics results, LD, and gene features directly from the knowledgebase.

The GenomicsDB also provides a genome browser, powered by *IGV.js*, which is an embeddable interactive JavaScript genome visualization tool developed by the Interactive Genomics Viewer (IGV) team [49]. For the GenomicsDB project, customizations have been made to allow *IGV.js* to query track data directly from underlying database, again using the WDK REST services, and from the NIAGADS Functional genomics repository (FILER), which provides harmonized functional genomics datasets that have been GIGGLE indexed for quick lookups [50,51]. Additional customizations provide links to NIAGADS gene and variant records in track-feature popups and coloring of features by NIAGADS annotations (e.g., ADSP annotations of variants, risk-association statistics) in track displays.

The browser is paired with a track selection tool to help users filter this extensive listing to find tracks of interest based on assay types and annotations captured in the track metadata. Like the summary statistics datasets, track metadata has been standardized, with biosample types, in particular, consistently annotated from the Uberon multi-species anatomy ontology (UBERON) [52,53], Cell ontology [54], and Cell Line Ontology [55].

## 3. Results

As of April 2023, the NIAGADS GenomicsDB offers access to summary statistics *p*-values from >80 GWAS and ADSP meta-analyses on both the GRCh37/hg19 and GRCh38/hg38 reference genome builds. Summary statistics are linked to >150 million ADSP annotated single-nucleotide variants and indels. Annotated variants in the GRCh37/hg19 version of the NIAGADS GenomicsDB include more than 29 million SNPs and approximately 50k short-indels identified during the ADSP Discovery Phase WGS and WES efforts [20]. The GRCh38/hg38 GenomicsDB builds on this to include additional variants from the ADSP’s ongoing efforts, with >232 million SNPs and short-indels identified from joint-genotype calling of 16,906 whole-genomes (R3 17k WGS). Of these, approximately 290k have significant AD or ADRD-risk association (*p* ≤ 5e^-8^) as reported in at least one NIAGADS GWAS summary statistics dataset to date. ADSP variants are highlighted in variant and dataset reports on the GenomicsDB website. The AnnotatedVDB (see **section 2.2.4**) also allows lookups and generation of annotated variant reports (see **section 3.2.2**) for variants of interest to the user that are present in dbSNP but currently lack genetic evidence associating them with AD/ADRD pathologies.

The standardization and harmonization effort described in the methods not only facilitates integration of large-scale annotation resources, but also helps optimize the database for big-data queries. This lets the knowledgebase query across millions of records and return compiled reports in real-time. This is most evident in GenomicsDB gene reports, which extract the top risk-associated variants (*p* < 0.001) from the GWAS summary statistics datasets within ±100kb of each gene and provide a comprehensive listing of both these variants and relevant annotations needed to make inferences about their potential regulatory impact on the gene.

The NIAGADS Alzheimer’s GenomicsDB creates a public forum for sharing, discovery, and analysis of genetic evidence for Alzheimer’s disease that is made accessible via an interface designed for easy mastery by biological researchers, regardless of background. The knowledgebase compiles all available data concerning summary statistics datasets and genetic evidence linking AD/ADRD to genes and variants into organized and easily navigated reports. Bioinformaticians can visit dataset report pages to interactively mine GWAS summary statistics datasets or use LocusZoom views of GWAS loci to weigh association strength of individual variants against that of others in its LD block. Clinicians and molecular scientists can look up their favorite genes or variants to peruse detailed reports that summarize known AD/ADRD associations within a broader genomic or functional genomic context.

### 3.1 Finding variants, genes, and datasets

The GenomicsDB homepage and navigation menu contain a site search that allows users to quickly find variants, genes, and datasets of interest by identifier or keyword (**Fig. 2**). Also offered is an interactive dataset browser that provides a full listing of all summary statistics datasets currently available in the GenomicsDB (**Fig. 3**). Users can find datasets of interest by manipulating the table in several ways, including searching by keyword (**Fig. 3A**) or applying advanced filters to identify datasets with specific sample or study design characteristics, such as clinical phenotypes, population, genotype, and sequencing center (**Fig. 3B**).

**Figure 2.**
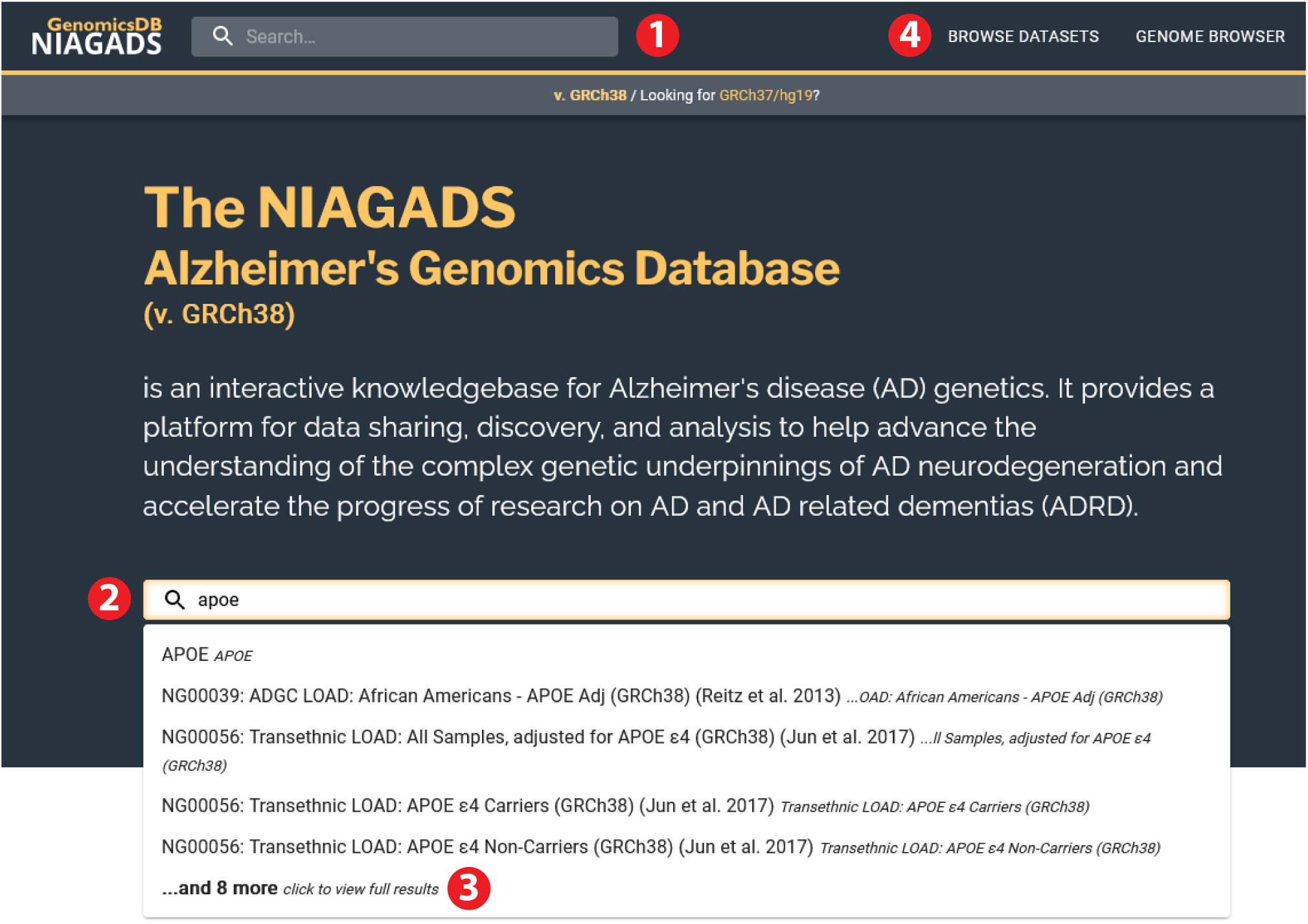
Searching the GenomicsDB. Search bars on the NIAGADS GenomicsDB navigation menu (1) and home page (2) allow users to quickly find a sequence feature using standard identifiers (e.g., genes: Ensembl, NCBI Entrez, official symbol; variants: refSNP identifier, chr:pos:ref:alt) or to perform a keyword search for genes and summary statistics datasets. Illustrated here are results found when searching for the gene “APOE”. Clicking on one of the top suggested search results will take the user to a detailed report. Users can also browse a listing of the full search results, which are reported as ranked lists of matching genes, variants, or datasets (3). A full listing of summary statistics datasets available in the GenomicsDB is accessed by selecting “Browse Datasets” from the main navigation menu (4). More details on the dataset browser are provided in Figure 3.

**Figure 3.**
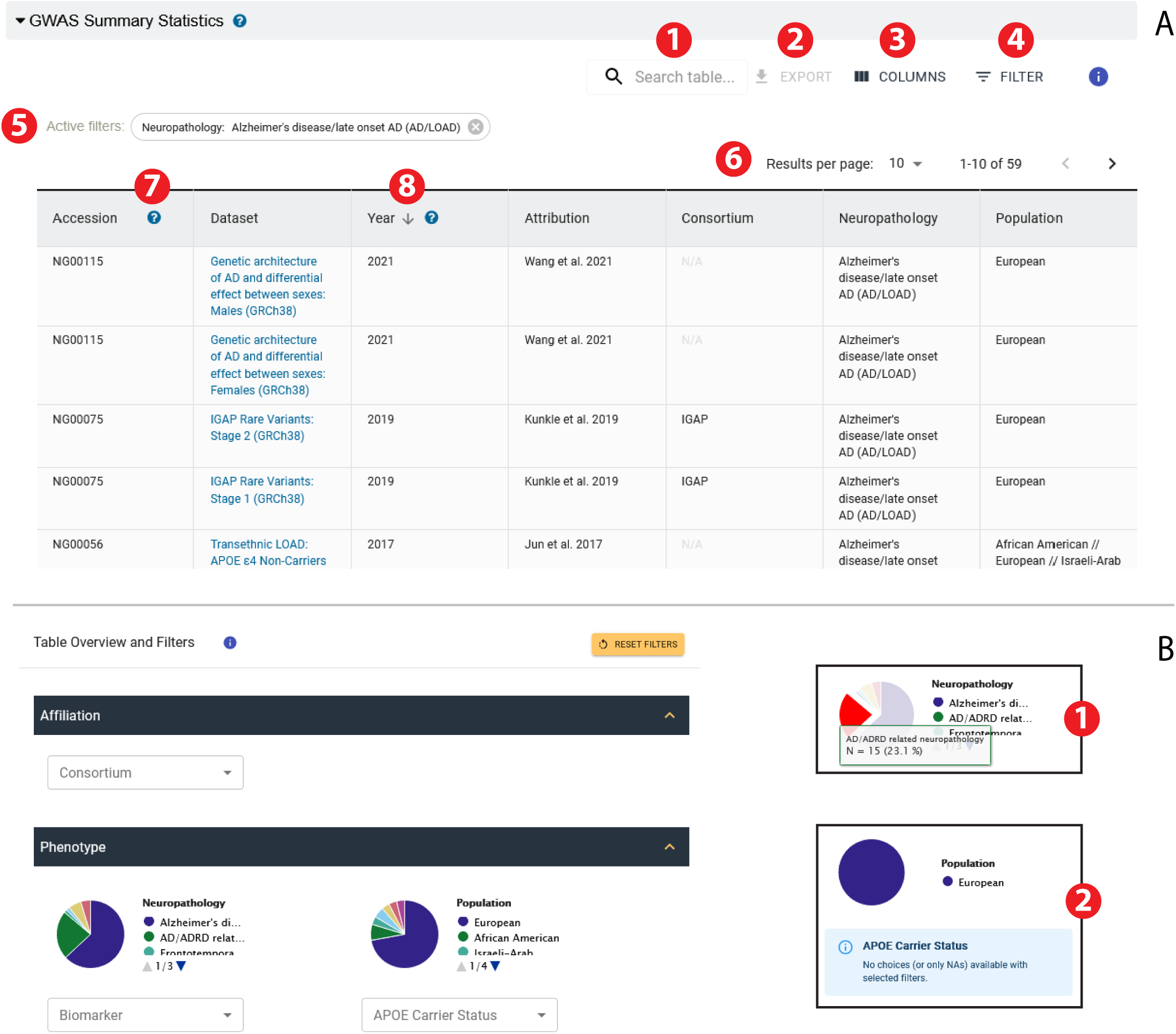
The GenomicsDB Dataset Browser provides a complete listing of summary statistics and meta-analysis results available in the resource. **A.** The browser can be searched by keyword (1) or mined using advanced filters that summarize phenotypes associated with sample cohorts (4; see also **3B**). Active advanced filters are listed above the table (5). Columns can be added or removed from the table (3) or sorted by contents (8) and sorted, modified, and/or filtered table exported as tab-delimited text (2). Table contents are paged to improve the site performance, but users can adjust the number of displayed rows (6). Help icons provide additional information about table fields (7). All tables in the GenomicsDB have similar functionality. **B.** The advanced filter interface summarizes the information in the table, providing drop down lists or interactive graphics to help guide users in data discovery. Hovering over chart elements or plot legends provides additional descriptive information and counts of table rows that match the filter criteria (inset 1). Selecting chart elements will apply the filter (i.e., the red pie slice in inset 1 indicates that the filter for AD-related neuropathologies was applied). Filter choices where update dynamically as new filter criteria are applied and alter the table contents (inset 2).

### 3.2 Browsing and mining reported data

Once a user identifies a gene, variant, or dataset of interest they are taken to a detailed report that summarizes all annotations in the GenomicsDB related to the search target and provides links to related features or datasets. GenomicsDB reports all have a standard layout, as illustrated in **Figures 4–5**. Reports contain a header that provides identifying and descriptive information and a graphical overview of the AD/AD-RD risk-associations in the dataset or linked to the genomic feature (**Fig. 4A,5A**). Headers also include link outs to the relevant third-party reference database for genes (Ensembl) and variants (dbSNP) and to the accession in NIAGADS for summary statistics datasets.

**Figure 4.**
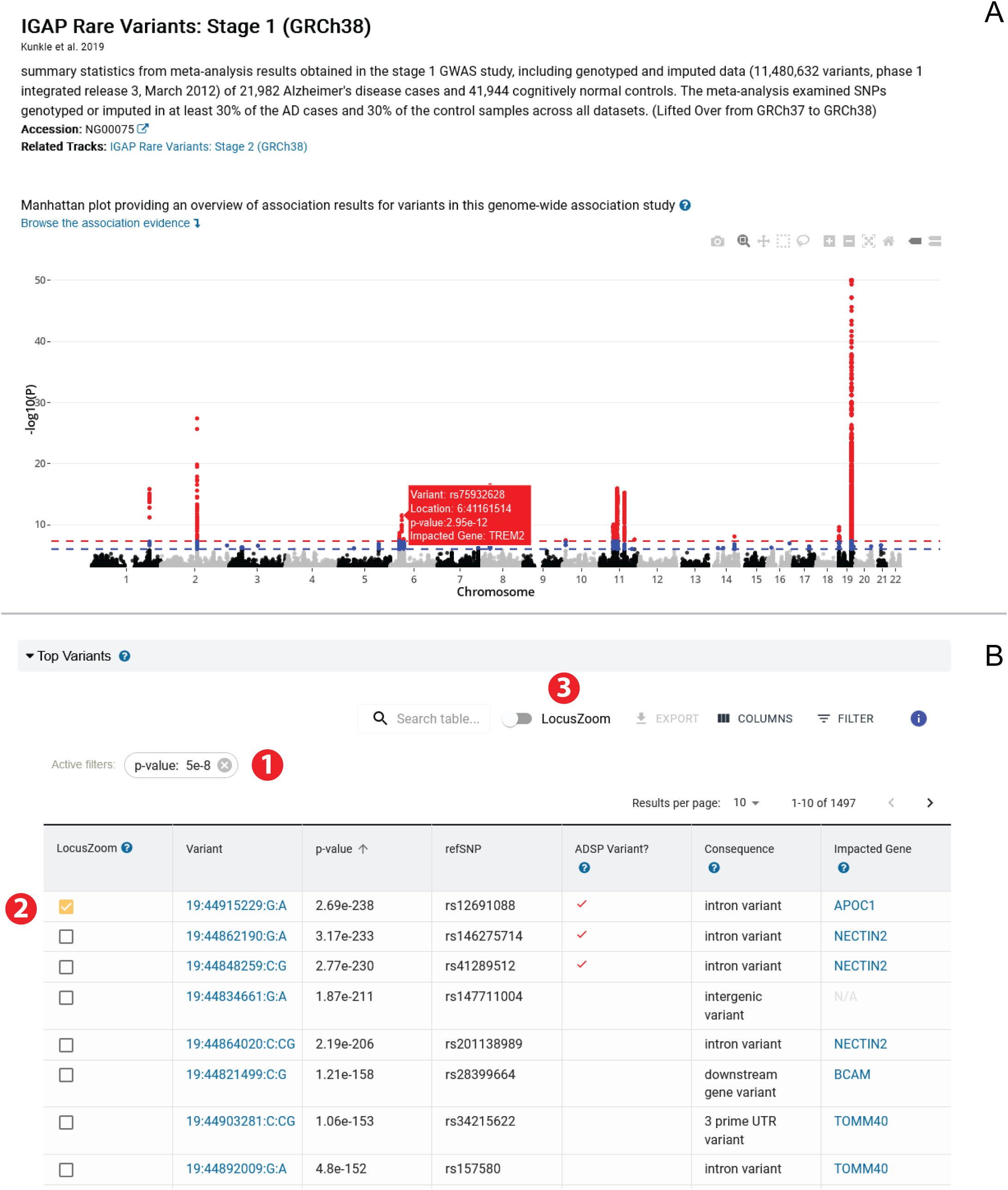
Example sections from a GenomicsDB dataset report. **A.** Dataset report header provides a summary, including an interactive Manhattan plot providing an overview of the dataset, links to related datasets in the GenomicsDB and to the NIAGADS repository for more information and instructions on making data access requests. **B.** The Manhattan plot is paired with a table found further down the page that lists the top hits (*p* ≤ 0.001) in the dataset, pre-filtered for variants meeting a genome-wide significance cut-off (p ≤ 5e^-8^) (1). The paired table can be is linked to the “Browse the association evidence” link in the page header to enable quick access. The table of top hits includes links to the variant reports (and impacted gene, when relevant) associated with each hit and is in turn, paired with LocusZoom (3) to allow users to select a variant (2) and view its association strength in its LD context.

**Figure 5.**
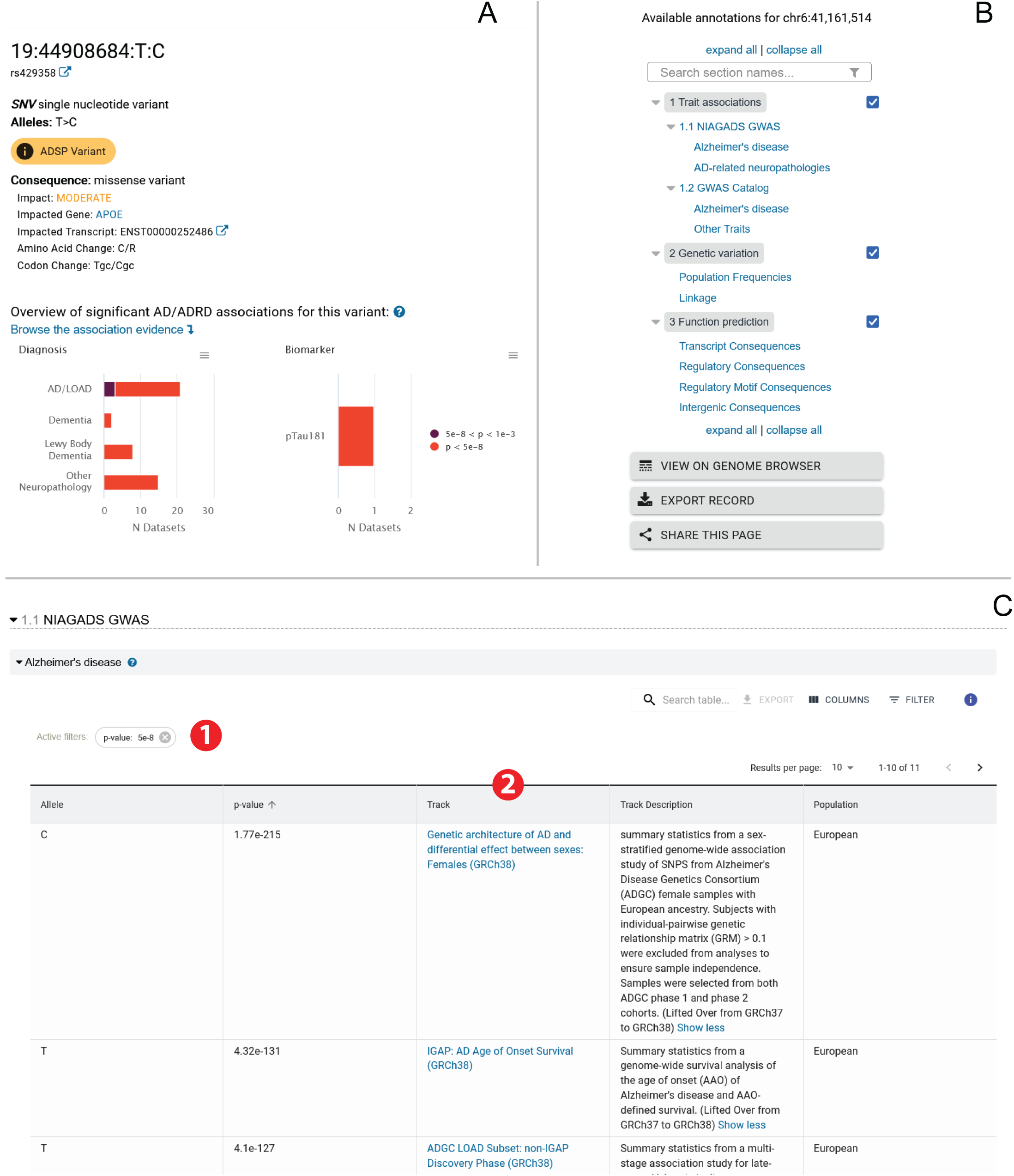
Example sections from a GenomicsDB Variant report. **A.** Variant report header provides summary information about the variant, including top predicted consequence from applying the ADSP annotation pipeline, whether the variant was included in an ADSP release (and passed quality control filters), quick links to alternative alleles or co-located variants, and a graphical summary of significant AD/ADRD risk-associations found in NIAGADS summary statistics datasets; comprehensive association results for the variant are listed in tables later in the report. **B.** Example navigation panel listing the structured information available in a variant report. Users can navigate directly to sections by clicking on the relevant link or toggle section visibility in the report using the checkboxes to the right. Also provided are buttons for exporting and sharing the report and for viewing the feature on the NIAGADS genome browser. **C.** Table of association results in NIAGADS AD GWAS summary statistics datasets for the variant highlighted in **A**. All top hits for the variant are report (p ≤0.001); the table is pre-filtered for associations meeting a genome-wide significance cut-off (p ≤ 5e^-8^) (1). Links are provided to the GenomicsDB reports for the relevant datasets (2).

All reports also contain a navigation panel (**Fig. 5B**) containing at a minimum a table of contents to help navigate the available information contained in the reports. Clicking on a report section in the navigation panel will scroll the page to the selected section. Navigation panels also include action buttons for sharing and exporting the information in the report, as well as one that loads a feature locus on the NIAGADS genome browser. Most data in GenomicsDB are presented in interactive tables that can be modified, exported, and filtered by keyword or more advanced filters (see **Fig. 3** for details).

#### 3.2.1 Summary statistics dataset reports

A comprehensive report is provided for each of the GWAS summary statistics and ADSP meta-analysis datasets in the NIAGADS GenomicsDB (**Fig. 4**). These reports allow users to browse the top risk-associated variants in the dataset and quickly isolate the genetic variants with genome-wide significance in the dataset (*p* ≤ 5e^-8^) via tables and interactive plots. The report header links back to the parent accession in NIAGADS where users can view the study details and publications, download the complete (genome-wide) *p*-values or make formal data access requests for the full summary statistics, related GWAS, expression, or sequencing data associated with the accession.

Dataset reports include an interactive Manhattan plot illustrating the distribution of risk-associated variants across the genome. Mousing over displayed points will display variant information (e.g., identifier, p-value, predicted impacted gene). Users can zoom into regions of interest and take snap shots of customized views. Static, high-resolution Manhattan plots are available for download.

The Manhattan plot is supplemented with a table that lists the top hits (*p*-value ≤ 0.001) in the dataset (**Fig. 4B**). This table is by default filtered for variants with genome-wide significance (*p* ≤ 5e^-8^) but the filter threshold can be adjusted as desired using the advanced filter tool. Also reported are predicted variant consequences that provide insight into the potential functional or regulatory impacts of the top variants (and proximal gene-loci); these annotations can be used to filter the table to help elucidate variants more likely to have a causal impact. All genes and variants listed in a dataset report are linked to corresponding feature reports in the GenomicsDB that offer detailed information about genetic evidence for AD/ADRD for the feature (see next sections). The table is paired with an interactive LocusZoom plot; the observed span and LD-reference variant can be changed by manipulating the plot directly or by selecting a row in the top hits table.

The GenomicsDB also generates reports for meta-analysis summary statistics generated by the ADSP that offer genetic evidence for gene-level and single-variant risk associations for AD. These are currently only available for the GRCh37/hg39 reference genome and result from a whole-exome sequencing (WES) case/control association analysis spanning 24 cohorts provided by the ADGC and the Cohorts for Heart and Aging Research in Genomic Epidemiology (CHARGE) completed as part of the ADSP discovery phase (NIAGADS ACCN:NG00065) [5].

#### 3.2.2 Variant reports

Variant reports include a basic summary about the variant (alleles, variant type, flanking sequence, genomic location) and a graphical overview of NIAGADS GWAS summary statistics datasets in which the variant has genome-wide significance (**Fig. 5A**). All other information in the report is subdivided into multiple sections that can be expanded or hidden at the user’s discretion (see navigation panel, **Fig. 5B**). These include selections on genetic variation (e.g., allele population frequencies and LD), function prediction determined via the ADSP annotation pipeline (incl. transcript and regulatory consequences), and comprehensive listings of GWAS inferred disease or trait associations from both NIAGADS summary statistics and the NHGRI-EBI GWAS Catalog (see **Fig. 5C** for an example). Tables listing summary statistics results can be dynamically filtered by *p-*value, dataset, phenotypes, or covariates, and the filtered results are downloadable. Links to the source datasets for each reported statistic are also provided, leading to detailed dataset reports (e.g., NIAGADS GWAS summary statistics) or to the source publication (e.g., curated variant catalogs). A table is also provided linking the GenomicsDB record to external variant annotation resources such as gnomAD, ClinVar (https://www.ncbi.nlm.nih.gov/clinvar/), and GTEx (eQTLs).

#### 3.2.3 Gene reports

GenomicsDB gene reports present the user with basic summary information about the gene (nomenclature, gene type, genomic span) and a graphical overview of NIAGADS GWAS summary statistics-linked variants proximal to or within the footprint of the gene that is paired with an interactive table listing the top risk-associated variants from the GWAS summary statistics datasets contained within ±100kb of the gene, similar in format to that reporting the top-hits for a summary statistics dataset (see **Fig. 4B**). Also provided on the gene report are sections reporting function prediction (GO associations and evidence) and pathway memberships. Links to external gene annotation resources (e.g., NCBI Gene, Ensembl, UniProtKB, the UCSC Genome Browser, and OMIM) are also provided.

### 3.3 Genome Browser

The NIAGADS GenomicsDB genome browser enables researchers to visually inspect and browse GenomicsDB data and annotations in a broader genomic context (**Fig. 6**). Tracks uniquely available on the NIAGADS genome browser include annotated ADSP variant tracks and tracks for each NIAGADS GWAS summary statistics dataset in the GenomicsDB, all of which are queried from the underlying database using REST services. Also available are >5000 functional genomics tracks, queried directly from FILER. To date these include data tracks from ENCODE, FANTOM5, GTEx, and the NIH Roadmap Epigenomics Mapping Consortium (Roadmap Epigenomics) [56].

**Figure 6.**
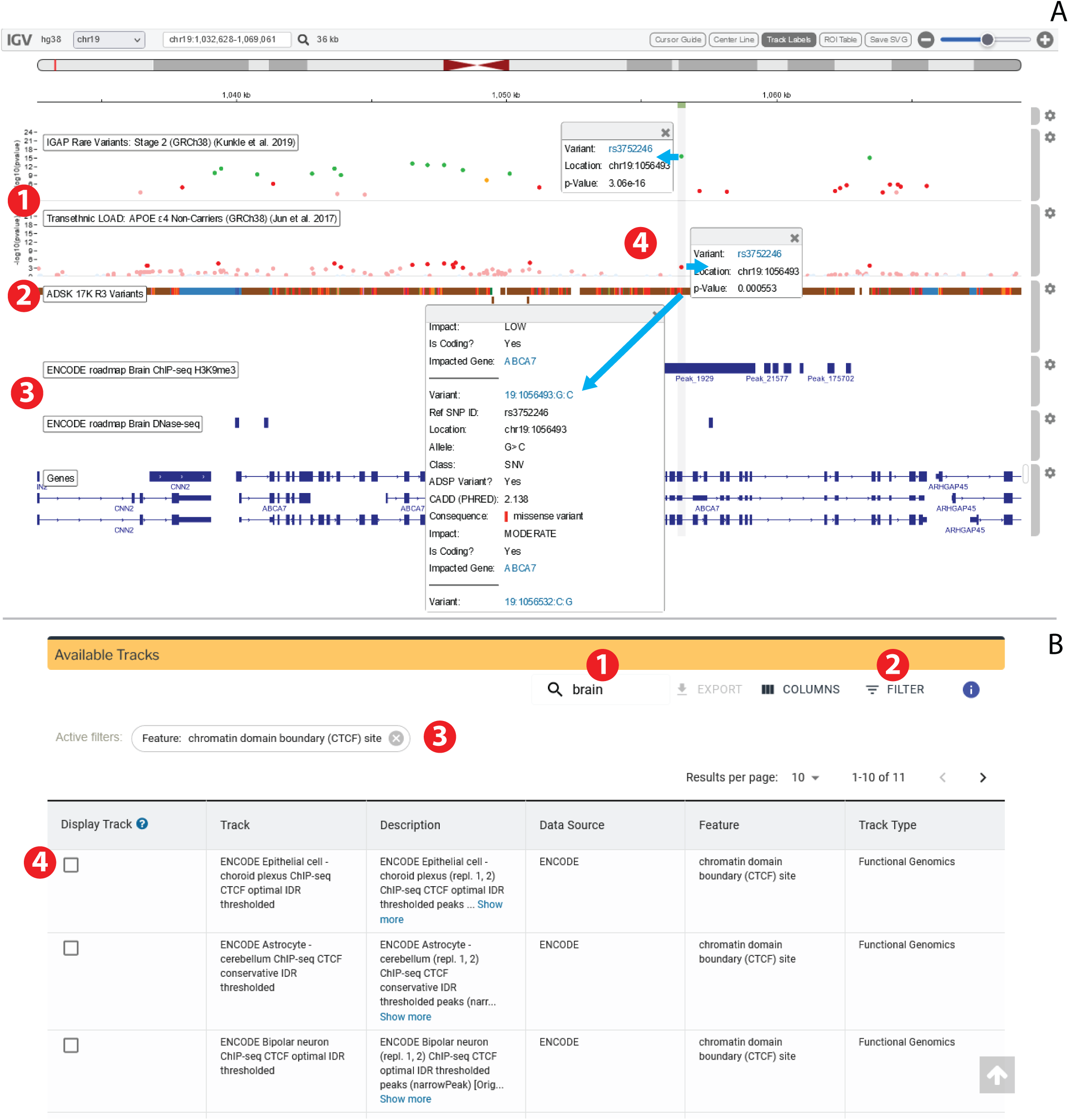
The NIAGADS Genome Browser is powered by *IGV.js* and allows users to visually inspect any of the NIAGADS GWAS summary statistics datasets in a broader genomic context. **A.** Example genome browser view illustrating a comparison between AD-relevant GWAS summary statistics datasets (1), annotated ADSP variants (2), and brain-related functional genomics tracks (3) in the region around the known AD-risk associated ABCA7 locus. Tracks are interactive; clicking on features will provide additional information (e.g., summary statistics or annotations) as well as a link to the selected gene or variant record (insets, 4). **B.** Interactive track selector allows users to browse all available tracks and search for tracks of interest by keyword, data source, biotype, functional genomics. The track listing can be searched by keyword (1) or using the advanced filters (2), with active filters listed above the table for easy removal (3). See Figure 3 for more details on manipulating and filtering GenomicsDB tables. Tracks can be added or removed from the browser by checking the row in the selector table (4).

Variant tracks are annotated by CADD scores and the most severe or deleterious predicted variant consequence as determined by the ADSP annotation pipeline. Annotations for individual variants can be viewed by clicking on the feature in the track or compared by adjusting the track coloring based on annotation valuee. Annotation track feature pop-ups also provide links back to the GenomicsDB variant records, as do those for GWAS summary statistics tracks (**Fig. 6A; highlight 4**). The reference gene track is colored by gene type; gene feature pop-ups provide details about the gene and its sub features and links back to the associated GenomicsDB gene record.

Users can discover tracks using the genome browser’s track selector (**Fig. 6B**). Tracks can be found via keyword search or by applying advanced filters that pull information from standardized track metadata, such as the original data source, biotype (e.g., cell or tissue), feature type (e.g., variant, gene, or regulatory element), or type of functional annotation. User tracks can be loaded via URL parameters; details about this functionality are provided in the site documentation.

## 4 Discussion

The NIAGADS Alzheimer’s Genomics Database is a user-friendly platform for interactive browsing and real-time in-depth mining of published genetic evidence and genetic risk-factors for AD. It provides unrestricted access to *p*-values from summary statistics of genome-wide association analysis of Alzheimer’s disease and related dementia or neuropathologies. Currently the GenomicsDB only disseminates summary statistics datasets pulled from the NIAGADS repository. However, future versions will include selected AD-relevant GWAS datasets (i.e., assessed for analysis strength, quality, relevancy, and study impact) pulled from the curated NHGRI-EBI GWAS Catalog and IEU OpenGWAS project.

Every entry in the GenomicsDB has been linked to relevant external resources and functional genomics annotations to supply further information and assist researchers in interpreting the potential functional or regulatory role of risk-associated variants and susceptibility loci. This is facilitated by a rigorous data standardization and harmonization process, which includes assigning genes and variants globally unique and persistent identifiers. This in turn creates opportunities for NIAGADS to facilitate further data sharing as it enables cross-references with external knowledge bases, as already established with the UniProtKB [57] and the Agora AD Knowledge Portal (doi:10.57718/agora-adknowledgeportal).

The GenomicsDB is a powerful research platform that provides a service for the AD genetics research community by both hosting comprehensive AD genetic and genomic findings and making them more accessible and creating opportunities for data reuse. It uses the latest web and database technologies to allow integration with new tools, and NIAGADS is constantly improving. The GenomicsDB front-end is updated periodically with enhanced features and new data visualizations. In addition, the REST services used to query the database to generate genome browser tracks or dataset and feature reports provide the foundation of an API that allows programmatic access to the database. Future work will focus on further developing this API to allow researchers to integrate standardized and harmonized GenomicsDB annotations in their own analysis pipelines. This work would allow users to retrieve variant annotations singularly or in bulk or pull out all harmonized annotations available for an arbitrary genomic span of interest.

The AD research community is actively encouraged through outreach and collaboration to submit data to NIAGADS to keep this public platform updated and timely. The database was designed to scale, with the rigorous data standardization practices and harmonized database underlying the GenomicsDB making it easy to add new data and annotations to the resource without requiring recalibration or reharmonization of the whole system with each new addition. As more data becomes available, we predict the NIAGADS Alzheimer’s Genomics Database will become a central hub for AD/ADRD research and data analysis.

## 6 Acknowledgements and Funding Information

This work is supported by the NIH National Institute on Aging with funding to the National Institute on Aging Genetics of Alzheimer’s Disease Data Storage Site (NIAGADS, U24AG041689), the Genome Center for Alzheimer’s Disease (GCAD), U54AG052427, and the Alzheimer’s Disease Genetics Consortium (ADGC) (U01 AG032984). Funding and acknowledgement statements for the ADSP can be found in the supplement.

